# Higher quality *de novo* genome assemblies from degraded museum specimens: a linked-read approach to museomics

**DOI:** 10.1101/716506

**Authors:** Jocelyn P. Colella, Anna Tigano, Matthew D. MacManes

## Abstract

High-throughput sequencing technologies are a proposed solution for accessing the molecular data in historic specimens. However, degraded DNA combined with the computational demands of short-read assemblies has posed significant laboratory and bioinformatics challenges. Linked-read or ‘synthetic long-read’ sequencing technologies, such as 10X Genomics, may provide a cost-effective alternative solution to assemble higher quality *de novo* genomes from degraded specimens. Here, we compare assembly quality (e.g., genome contiguity and completeness, presence of orthogroups) between four published genomes assembled from a single shotgun library and four deer mouse (*Peromyscus* spp.) genomes assembled using 10X Genomics technology. At a similar price-point, these approaches produce vastly different assemblies, with linked-read assemblies having overall higher quality, measured by larger N50 values and greater gene content. Although not without caveats, our results suggest that linked-read sequencing technologies may represent a viable option to build *de novo* genomes from historic museum specimens, which may prove particularly valuable for extinct, rare, or difficult to collect taxa.

## Introduction

A disconnect between the capabilities of high-throughput sequencing technologies and the quality, or lack thereof, of historic museum specimens has largely neutered the ability of genomic methods to access molecular data from degraded specimens. Natural history collections (NHCs) store a wide variety of species from across the globe, including those that are difficult to collect or extinct in the wild. Voucher specimens housed in NHCs have been an invaluable source of morphological material as they provide a reference for measuring change across both space and time. More recently, specimens contained in NHCs have been recognized as important repositories of genetic data (Payne & Sorenson, 2002; Wandeler, Hoeck & Keller, 2007) and have provided insight into the phylogenetic relationships and origins of species (Suarez & Tsutsui, 2004; McLean et al., 2015). Quick progress in genomics methods are now enabling the use of museum specimens in ways that were not imaginable until only a few years ago. “Museomics”, or the application of genomic techniques to museum specimens, has already uncovered reticulate evolutionary histories across hominids (Green et al. 2010; Meyer et al., 2012, 2014) and is increasingly resolving the phylogenetic Tree of Life (Teeling & Hedges, 2013; Lessa, Cook, D’Elia, & Opazo, 2014; Wood, González, Lloyd Coddington, & Scharff, 2018), with expanded applications including, but not limited to, identifying functional variants implicated in ecological adaptations (Opazo, Palma, Melo, & Lessa,. 2005) and estimating mutation rates and the timing of evolutionary events (Pélissié, Crossley, Cohen, & Schoville, 2018).

Over time, however, and through exposure to agents known to degrade nucleic acids (UV, temperature, pH, salt, chemical modification, etc.; Dessauer, Cole, & Hafner, 1990; Lindahl, 1993; Dean & Ballard, 2001; Willerslev & Cooper, 2005), DNA degrades into short fragments, which can complicate the application of genomic methods to museum specimens. Since the 1970s, when museums widely began archiving tissues, collection and preservation methods have varied widely, but generally evolved to accommodate changing analytical technologies, resulting in the variety of preservation methods (e.g., formalin, ethanol, ground, frozen, etc.) and quality of tissue collections available to researchers today. In addition to the challenges of tissue preservation, field conditions including weather, processing speed, and available cold storage options are inherently unpredictable, resulting in further inconsistencies in field-collected tissue quality.

*De novo* genome assembly is the computational process of optimally fitting short-read fragments output from sequencers into a larger contiguous whole-genome sequence, recovering critical information about the locations of genes and variants that are lost in the sequencing process. Assembly methods are based on the often-incorrect assumption that similar DNA fragments originate from the same position within the genome; therefore, assembly can be complicated by the presence of extended repeats or regions of high divergence that extend beyond the sequenced read length (Alkan, Sajjadian, & Eichler, 2011; Nagarajan & Pop, 2013). Unfortunately, methods that yield the highest quality *de novo* genome assemblies often require large quantities of high molecular weight (HMW) DNA as starting material for library preparation, as the ability to resolve sequencing artefacts in assembly improves with increasing read length. This prerequisite often makes these methods inaccessible to degraded specimens. For example, although the recent emergence of long-read sequencing technologies (>10-50 kb) has significantly improved the computational complexities of *de novo* genome assembly, long-read sequencing requires large quantities of HMW DNA as a starting material, making these methods impractical for most museum samples (Rowe et al., 2011). Prior to the development of long-read sequencing, the most common approach to *de novo* genome assembly has involved a combination of shotgun short-insert (<500 bp) and mate-pair long-insert (>2000 bp) libraries of varying insert size, where the first would be used for assembly and the second for scaffolding. Once again, scaffolding would be limited by fragmented DNA, as input molecules must be longer than the selected insert size. More recently, the protocol accompanying the assembler DISCOVAR denovo (Broad Institute, 2015; Weisenfeld, Kumar, Shah, Church, & Jaffe, 2017), which is based on single short-insert shotgun libraries sequenced to ∼60X using 250 bp paired-end reads, appears to be a viable option for genome assembly from degraded samples. This approach proved cost-effective for the genome assembly of 20 *Heliconius* species (Edelman et al., 2018), but for organisms with larger genomes this option is significantly more expensive than other approaches due to the high coverage and longer read lengths required. An appealing alternative is reference-guided assembly (Rowe et al., 2011; Staats et al., 2013), where either raw reads are mapped to an existing high-quality reference genome from a closely related species to build a consensus sequence (Pop, 2009) or a related reference genome is used only as a scaffolding guide (Gnerre, Lander, Lindblad-Toh, & Jaffe, 2009). While this approach may offer a partial solution, high quality, closely related references, a prerequisite for this approach, are not available for a large number of ecologically relevant taxa yet. To overcome this obstacle, other studies have recommended avoiding whole-genome sequencing (WGS) of museum specimens altogether, suggesting exome capture (Bi et al., 2013) or other reduced-representation approaches (Jones & Good, 2018) as an alternative proxy for accessing molecular data from museum specimens. However, in addition to complex laboratory work, these approaches retrieve only a restricted subset of sequence data relative to WGS. In addition to the potentially confounding effect of pervasive purifying selection on exonic coding regions (Jackson, Campos, & Zeng, 2014), exome sequencing further fails to represent significant regulatory or non-coding regions essential to phylogenetic reconstruction (Nei & Tateno, 1975; Lynch, 1989), and, with increasing awareness, for understanding the targets of adaptive evolution (Andolfatto, 2005; Brooks, Turkarslan, Beer, Lo, & Baliga, 2011).

In the grey area between second and third generation sequencing, linked-read or ‘synthetic long-read’ (SLR, Voskoboynik et al., 2013) sequencing may provide a cost-effective solution for *de novo* genome sequencing from degraded specimens. These methods allow the assembly of pseudo-long reads up to 18kb from short-read data with higher accuracy compared to true long-read sequencing techniques (Jiao & Schneeberger, 2017). Initially introduced by Illumina (Kuleshov et al., 2014; McCoy et al., 2014), SLR methods have not been widely adopted by evolutionary biologists and museum scientists. 10X Genomics (Zheng et al., 2016), a newer technology loosely based on innovations developed by the Illumina SLR technique, offers several advantages for museum science applications. Specifically, this method requires as little as 1 nanogram of input material and it is robust to the effects of input DNA quality. 10X Genomics uses microfluidics to split extracted DNA fragments across >100,000 partitions or ‘GEM’s (gel-coated beads). Each ‘GEM’ then contains a fraction (< 0.5%) of the genome, which is further sheared and barcoded. Reads from the same partition or ‘GEM’ are sequenced via conventional Illumina short-read sequencing and assembled locally, by barcode, as they must be derived from the same original DNA fragment (Goodwin, McPherson, & McCombie, 2016; van Dijk, Jaszczyszyn, Naquin, & Thermes, 2018). Although HMW DNA is optimal for any method, the physical separation of DNA fragments in a ‘GEM’ largely eliminates the issue of degraded DNA and increases assembly confidence by geographically linking small-reads in genome-space, thereby reducing misassembly. This library preparation method can also facilitate allele phasing and the detection of structural variants, although its power will depend on the quality of starting DNA (Lee et al., 2016; Zheng et al., 2016).

While linked-reads are currently optimized for human genomes (kb.10xgenomics.com) and most often applied to cancer and biomedical related questions (Zheng et al., 2016), these methods are beginning to be explored in other taxa (orchids, Zhang et al., 2017; sea otters and beluga whales, Jones et al., 2017a, b). As a consequence of phylogeny, 10X Genomics methods are most easily extended to other mammalian taxa, expected to have similar genome size and structure (*e.g.*, repeat content, heterozygosity, etc.). Thus, as a proof-of-concept, we compare assembly quality and content of four deer mouse (*Peromyscus*) genomes sequenced and assembled using the 10X Genomics linked-read approach with — at a comparable cost — four publically-available shotgun Illumina mammalian genome assemblies generated from comparable read volumes. We demonstrate the utility of this economical approach to whole genome reconstruction for researchers interested in questions related to systematics and functional genomics.

## Materials & Methods

Twenty-five micrograms of frozen liver tissue from each of four field-collected museum specimens (*Peromyscus attwateri* [MSB:Mamm:84733], *Peromyscus aztecus* [MSB:Mamm:48205], *Peromyscus melanophrys* [MSB:Mamm:273915], and *Peromyscus nudipes* [MSB:Mamm:70743]) were loaned from the Museum of Southwestern Biology (MSB). Three of the specimens were collected internationally and collections dates ranged from 1982 to 2006 (Table 1). Genomic DNA was extracted using a standard QIAGEN Genomic Tip (Valencia, CA, USA) protocol. DNA was quantified with Qubit and its quality was assessed using an Agilent TapeStation (Santa Clara, CA, USA). DNA from each of the four species contained a distribution of DNA fragments heavily skewed towards smaller molecular weights. As fragment size distribution greatly influences the contiguity of the genome assembly, we further processed the samples using the Circulomics short read eliminator kit (Baltimore, MD, USA), which removes DNA molecules shorter than 10kb, and progressively up to 25 kb, thereby removing the vast majority of our DNA sample. The remaining DNA from each of these samples was sent to the Genomics Core Facility at Icahn School of Medicine at Mount Sinai for library preparation, where samples were run on a Femto Pulse (Agilent, Santa Clara, CA, USA) to assess fragment size distribution post-Circulomics (Supplemental Information). Resulting 10X libraries were sequenced at Novogene (Novogene, Sacramento, CA, USA) using 150 bp paired reads generated in one lane of Illumina HiSeq X for each species. Raw data were assembled with Supernova v. 2.1.1 (Weisenfeld *et al.* 2017) and the final fasta file was generated using the ‘pseudohap style’ option in *supernova mkoutput* using default settings. All commands used for this work are available at https://github.com/macmanes-lab/museum_genomics.

**Table 1.**
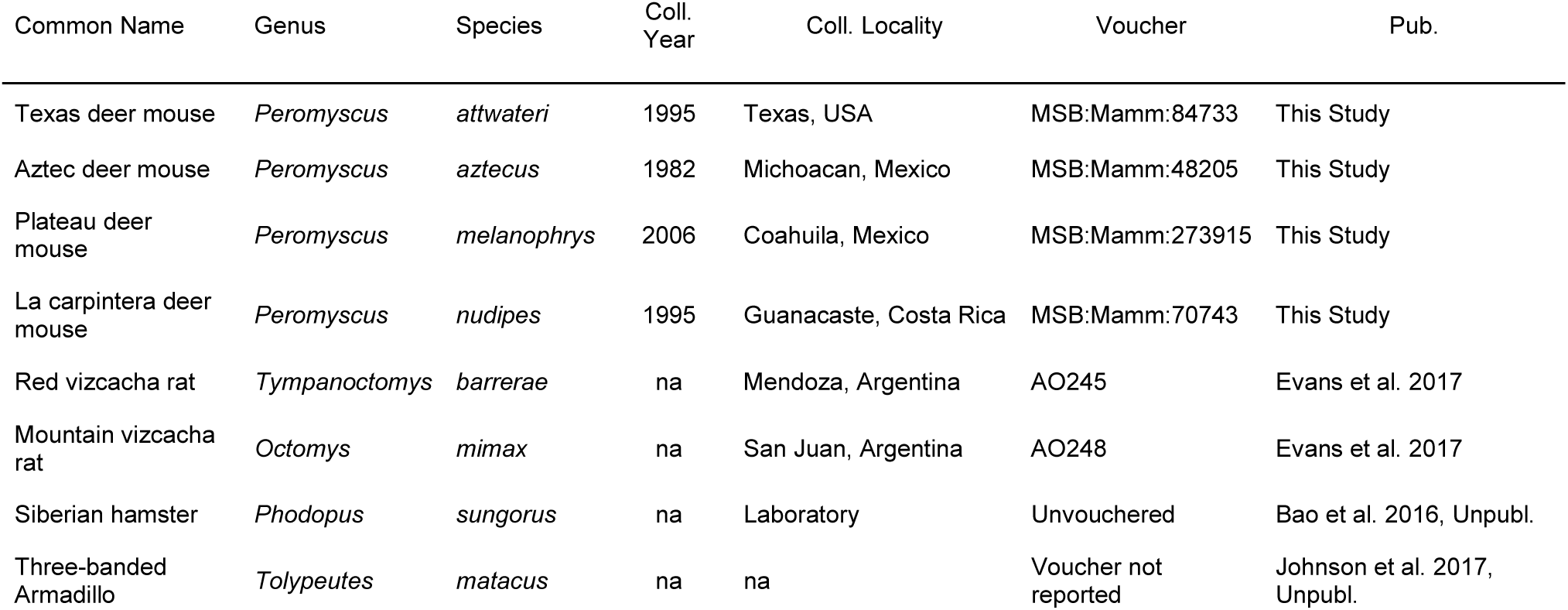
Natural history data for specimens sequenced using 10X Genomics (*Peromyscus* spp.) and publically available single-land Illumina assemblies. Coll. = collection. Pub. = publication.

We downloaded four publically available genome assemblies generated from a single shotgun library sequenced on an Illumina platform. To minimize differences in genome structure that could affect the performance of different methods, we selected four mammalian species of similar genome size including *Tympanoctomys barrerae, Octomys mimax* (Evans, Upham, Golding, Ojeda, & Ojeda, 2017), *Phodopus sungorus* (Bao, Hazelerigg, Prendergast, & Stevenson, 2016), and *Tolypeutes matacus* (Johnson et al., 2017).

To compare 10X versus shotgun-based assemblies, we assessed genome quality through comparison of relative N50 values, genome completeness using presence of BUSCO genes and of orthologous groups. Because the mammalian genomes considered here are generally similar in size, N50 values are comparable and normalization by genome size is not necessary. Comparative shotgun assemblies were selected based on the number of total reads sequenced (∼200M), as an equivalent sequencing cost comparison against 10X Genomics. Read counts and assembly details for each externally sourced genome are available in Table 2.

**Table 2.**
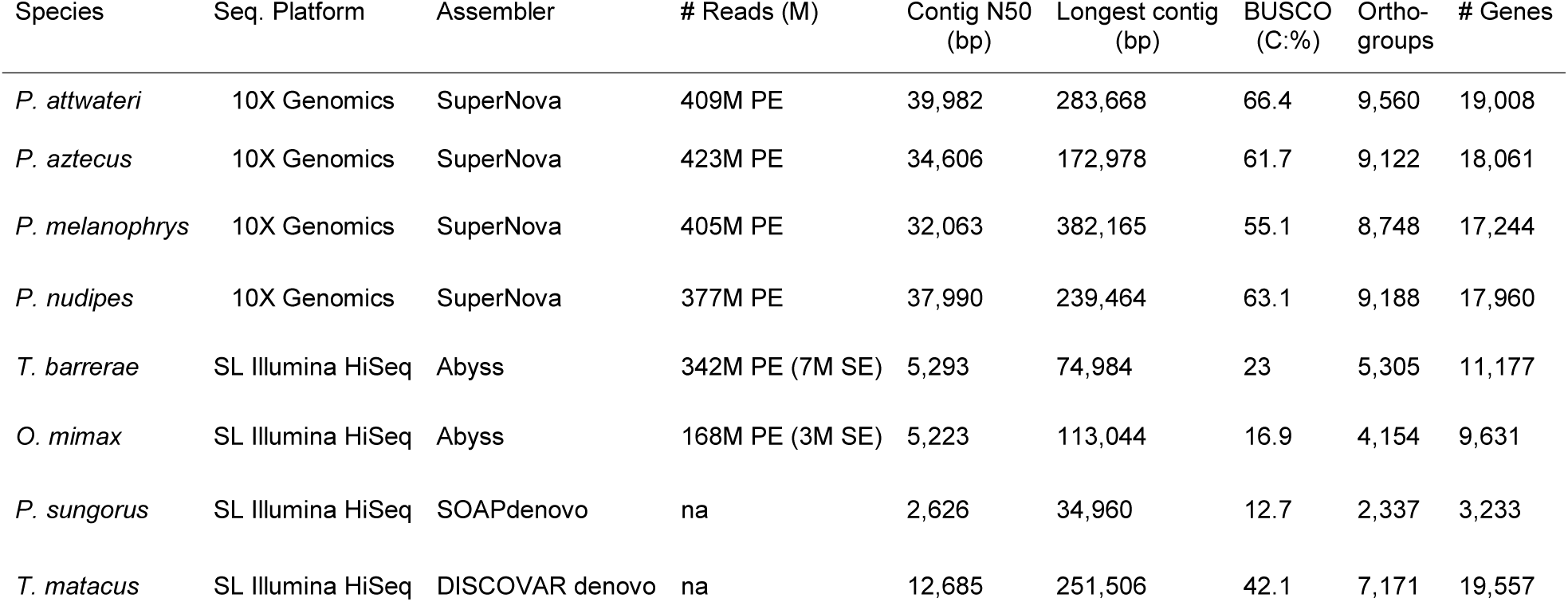
Sequencing and assembly quality statistics for each examined genome, including: Sequencing Platform (Seq. Platform), Assembler, Number of Reads (# of Reads [M = million], PE [paired-end], SE [single-end]), Contig N50 in base pairs (bp), longest contig (bp), percent (%) complete (C) BUSCOs, and the number of genes (# Genes) annotated.

All genomes were annotated using Maker v. 2.3.1 (Cantarel et al., 2008) using the *Mus musculus* (GCF_000001635.26) reference proteome. The N50 statistic was calculated via the *abyss-fa*c tool included in the Abyss package (Simspon et al., 2009). Benchmarking Universal Single-Copy Orthologs (BUSCO V. 3; Simão, Waterhouse, Ioannidis, Kriventseva, & Zdobnov, 2015) statistics were used as metrics of genome completeness based on gene content for genes conserved across Mammalia (mammalia_odb9).

We grouped genes from each species into orthogroups using OrthoFinder v. 2.3.3 (Emms & Kelly, 2015) and determined the number of orthogroups we could retrieve from each assembly. As proof of concept, a species tree was built for the *Peromyscus* taxa sequenced here, three publically available *Peromyscus* genomes (*P. maniculatus* [GCA_003704035], *P. leucopus* [GCF_004664715], and *P. polionotus* [GCA_003704135]), and four outgroup sequences (*Rattus norvegicus* [GCA_000001895.4], *Mus musculus* [GCF_000001635.26_GRCm38], *Onychomys torridus* [GCA_004026725], and *Sigmodon hispidus* [GCA_004025045] based on the OrthoFinder sets of orthologous genes, using IQTREE and default settings (Nguyen, Schmidt, von Haeseler, & Minh, 2015).

## Results

Read counts for each analyzed genome are available in Table 2. Additional assembly statistic (n:500, L50, N80, N50, N20, E-size, etc.) are available online at https://github.com/macmanes-lab/museum_genomics/blob/master/assembly_stats.md.

N50 values ranged from 2,626 to 39,982 bp with the highest values for 10X Genomics assemblies (36,160 bp on average) and lowest for single-lane Illumina assemblies (6,457 bp on average; Table 2). The number of genes annotated ranged from 3,233 (*P. sungorus*) to 19,008 (*P. attwateri*), with 10X Genomics assemblies 18,068 genes on average, compared to 10,900 genes in the average shotgun-based assembly. BUSCO measures of genome completeness ranged from 16.9% (*O. mimax)* to 66.4% (*P. attwateri*) and were again highest for 10X Genomics assemblies (average: 61.6%) and lowest for shotgun assemblies (average: 27.3%; Table 2).

Annotations and predicted transcripts and proteins are available at http://doi.org/10.5281/zenodo.3351485. OrthoFinder identified 5,305 orthologous groups on average in shotgun-based assemblies and 9,112 on average in 10X assemblies (p < 0.05; Table 2). Our basic maximum-likelihood species tree resolved relationships with 100% bootstrap support (Fig. 1). Raw reads and assemblies are available through The European Nucleotide Archive (ENA) under project number PRJEB33530.

**Figure 1.**
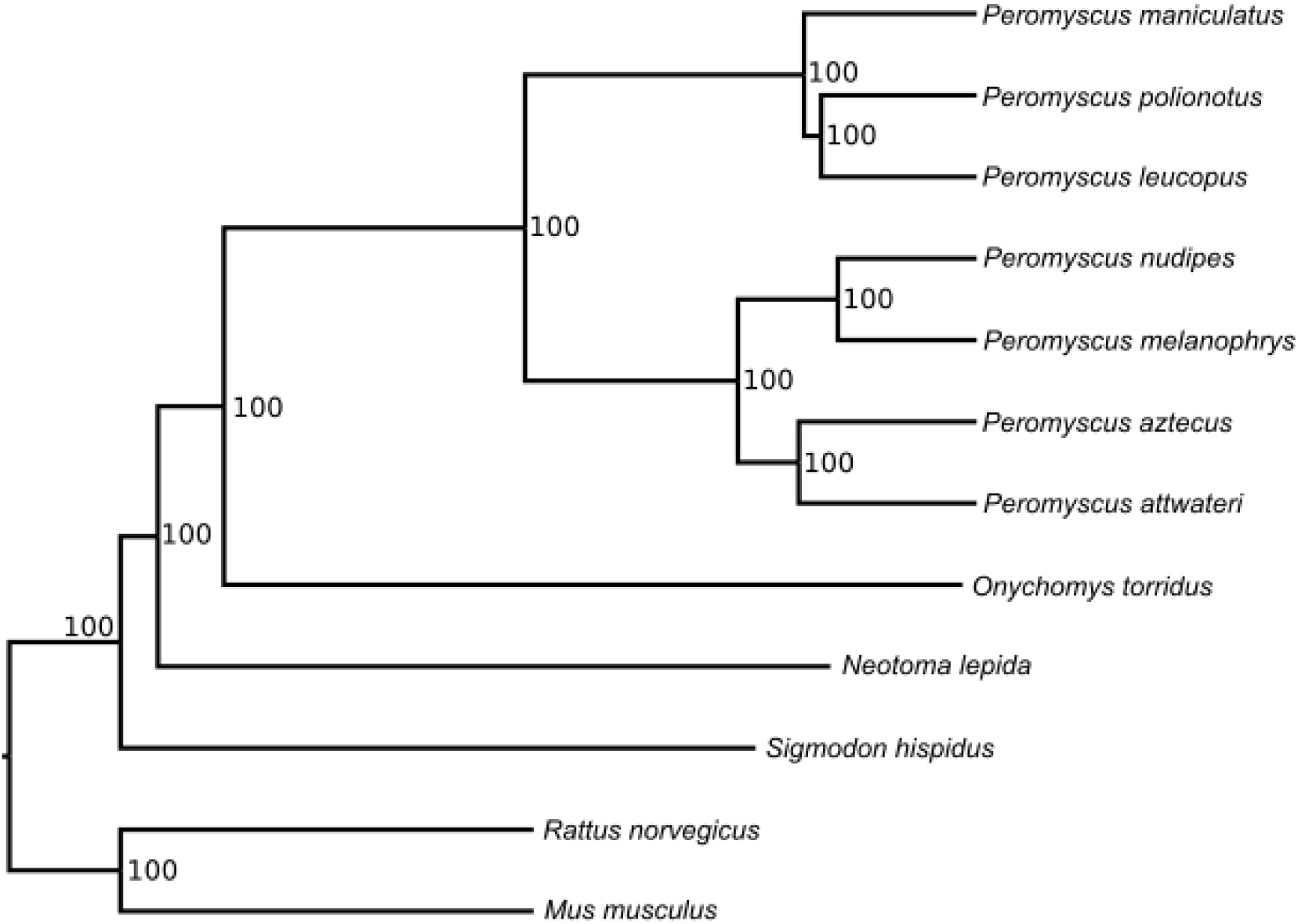
Maximum-likelihood phylogeny of examined *Peromyscus* species generated from consensus orthogroups, demonstrates complete resolution (100 bootstrap support for all nodes).

## Discussion

Linked-read sequencing facilitates the production of higher quality *de novo* genomes from historic samples, in less time, and with less effort than traditional shotgun based methods, providing a new option for accessing the genomes of aged samples. As such, linked-read sequencing may be the long-awaited key to unlocking the molecular secrets of NHCs and have applications across a broad range of evolutionary and ecological questions. *De novo* assemblies from linked-reads have greater contiguity and completeness relative to *de novo* assemblies based on shotgun libraries for comparable read volumes (Table 2). With the same sequencing effort (*e.g.*, 200 million 150 bp paired-end reads) linked-read sequencing results in a six-fold increase in N50 values and increases the number of represented genes by three times, without requiring a reference sequence from a closely related species. Using 10X assemblies as reference genomes to map population-level whole genome resequencing data, which are only minimally affected by DNA degradation, will also increase the amount of variation, both sequence and structural, available for genotyping within and among species. Note that, once a *de novo* reference genome is available, shotgun libraries sequenced with short reads will still be a viable and cost-cost effective methodological choice for whole-genome resequencing. Linked-read methods facilitate the detection of rare alleles and enable haplotype phasing, both of which may be key to identifying emerging model taxa for biomedical research, investigations of rare genetic disorders in humans, and analyses of introgression, by enabling estimates of local ancestry (Tennessen et al., 2012; Janzen, Wang, & Hufford, 2019). Previously limited by technology, molecular investigations of museum specimens traditionally centered around systematic inquiry and phylogenetics. Now, the ability economically generate quality *de novo* assemblies for lower-quality tissue resources increases the power of these historic archives to address new questions.

Although the quality and completeness of linked-read assemblies are still dependent on DNA integrity, the application of linked-read methods may be especially impactful for rare or extinct species or when the collection of new material is difficult or impossible (Payne & Sorenson, 2002) due to the conservation status or geographic location (*e.g.*, international) of the target species. As new or higher-quality tissue samples will never again be available for extinct species, linked-reads offer an improved method for accessing data from preserved tissues of these species, even if the generation of perfectly contiguous genomes for these taxa is not attainable. In cases where the target species is highly divergent from available reference sequences, such as the case for extinct species or otherwise exceptionally divergent taxa (*e.g.*, monotypic genera [*Ailurus, Eira*] or families [*Dugongidae, Orycteropodidae*]), *de novo* genome assemblies, rather than reference-based assemblies may provide more information.

As a caveat, 10X Genomics methods have not yet been tested for genomes larger than ∼3Gb (e.g., human-sized), so although they are appropriate for many mammalian species, they may be less applicable to species with larger genomes. Detailed analysis of structural variation, as is often implicated in ecological adaptation (Wellenreuther, Mérot, Berdan, & Bernatchez, 2019), remains under the purview of long-read or hybrid (short and long reads) *de novo* sequencing methods and N50 statistics for linked-read assemblies are still limited relative to true long-read methods. Although the number of raw reads are variable within both groups — the shotgun-based and 10X Genomic assemblies — the number of reads is not correlated with genome quality. This leads us to conclude that differences in assembly quality are not driven by differences in sequencing depth. Finally, although our results are derived from lower quality frozen tissue samples, tissues remain unavailable for many pre-molecular era specimens. While linked-reads may be a solution to produce a *de novo* genome from poor quality tissues, this method has not been applied to and may not be appropriate for highly degraded museum study skins or destructively-sampled bone remains. Reduced-representation genetic approaches (Sanger sequencing, RADseq) or enriched sequencing methods (Bi et al., 2013; Staats et al., 2013; Jones & Good, 2016) may remain the most effective means of extracting data from more historic specimens in the absence of a closely related reference genome.

Ultimately, our results underscore the importance of continued scientific collecting and the archival of personal legacy collections into NHCs into the future, as new technologies will continue to improve our ability to extract molecular information from degraded and aged samples. The centralization of biological resources and associated information ensures the broad utility of these specimens to the scientific community and facilitates tests, such as these, to determine the best available means of extracting meaningful sequence data from lower quality DNA. In particular, we endorse maximizing the utility of a specimen through the archival of multiple tissue types, through multiple storage media (liquid nitrogen, ethanol, RNA later® [Sigma-Aldrich, St. Louis, Missouri, USA], etc.) to maximize future uses of these archives as technologies continue to evolve (Lessa et al., 2014; McLean et al., 2015).The ability to generate WGS data from field-preserved tissues, further encourages the expansion of resurvey projects (such as the NSF funded Grinnell Resurvey Project from the Museum of Vertebrate Zoology, UC Berkeley) as a means for measuring change through time (Moritz, Patton, Conroy, Parra, White, & Beissinger, 2008) and opens the possibility of sequencing *de novo* genomes from now extinct species with preserved tissues available. In an era of unprecedented ecological and environmental change (Ceballos, Ehrlich, & Dirzo, 2017), genomic analyses of historic samples will help us understand the evolutionary responses of natural populations to environmental perturbation and hence lay the foundation for proactive management initiatives and predicting future responses (Wandeler et al., 2007; Malaney & Cook, 2013).

## Acknowledgements

We thank the Museum of Southwestern Biology for the *Peromyscus* tissue loans that made this work possible and the National Institute of Health (Grant Number: NIH R35GM128843) for funding.

## Data Accessibility

All commands used for this work are available at https://github.com/macmanes-lab/museum_genomics. Raw reads and assemblies are available through The European Nucleotide Archive (ENA) under project number PRJEB33530, with assembly IDs: *P. attwateri* (ERZ1029326), *P. nudipes* (ERZ1029275), *P. melanophrys* (ERZ1029325), *and P. aztecus* (ERZ1029324).Assembly statistics are available online at https://github.com/macmanes-lab/museum_genomics/blob/master/assembly_stats.md. Annotations and predicted transcripts and proteins are available at http://doi.org/10.5281/zenodo.3351485.

## Author Contributions

All authors contributed equally to this manuscript. MDM assembled the *Peromyscus* genomes.

