## Supplemental Information for "Higher quality *de novo* genome assemblies from degraded museum specimens: a linked-read approach to museomics"

Distribution of fragment size (bp) and relative fluorescence units (RFU) for *Peromyscus aztecus.*

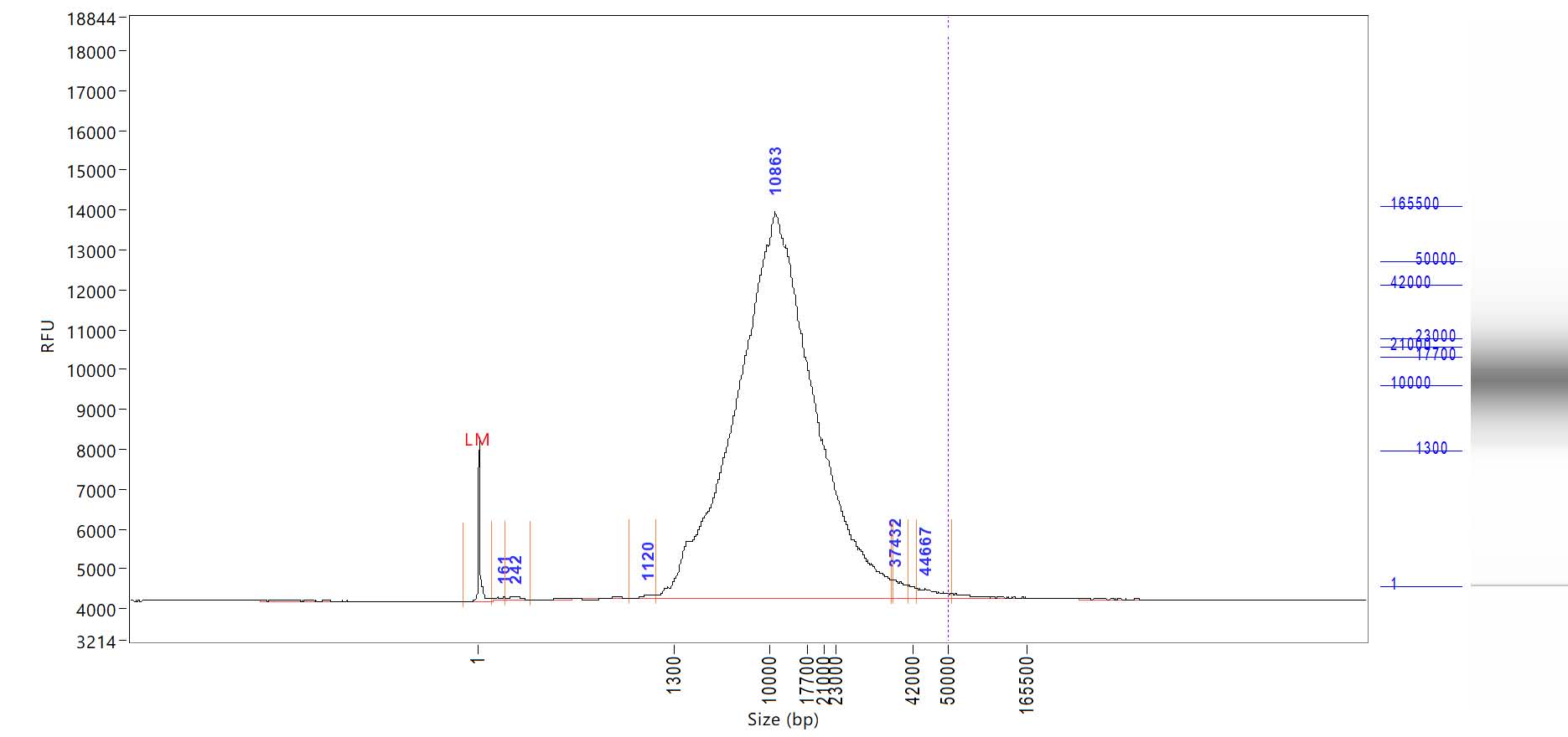

Distribution of fragment size (bp) and relative fluorescence units (RFU) for *Peromyscus melanophrys.*

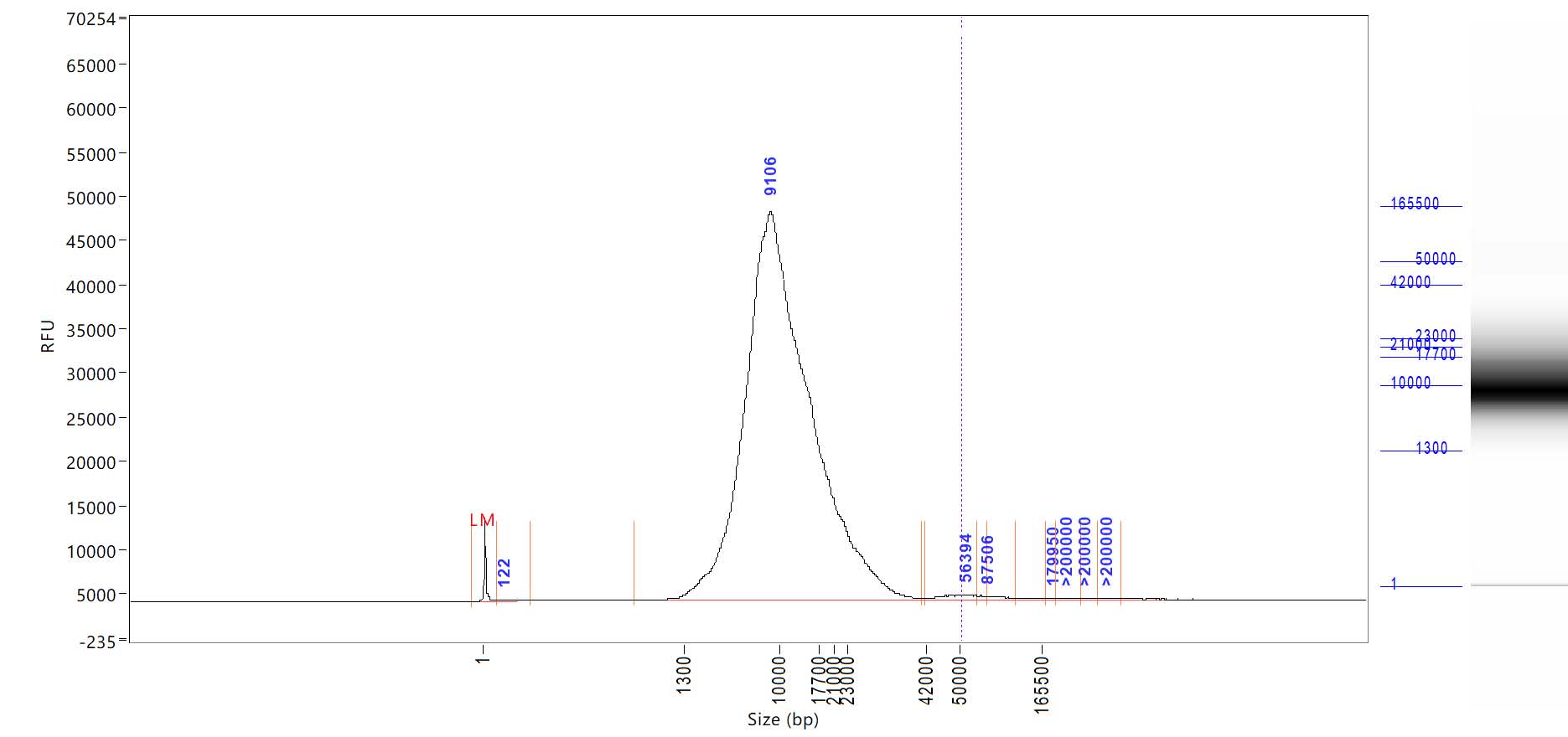

Distribution of fragment size (bp) and relative fluorescence units (RFU) for *Peromyscus nudipes.*

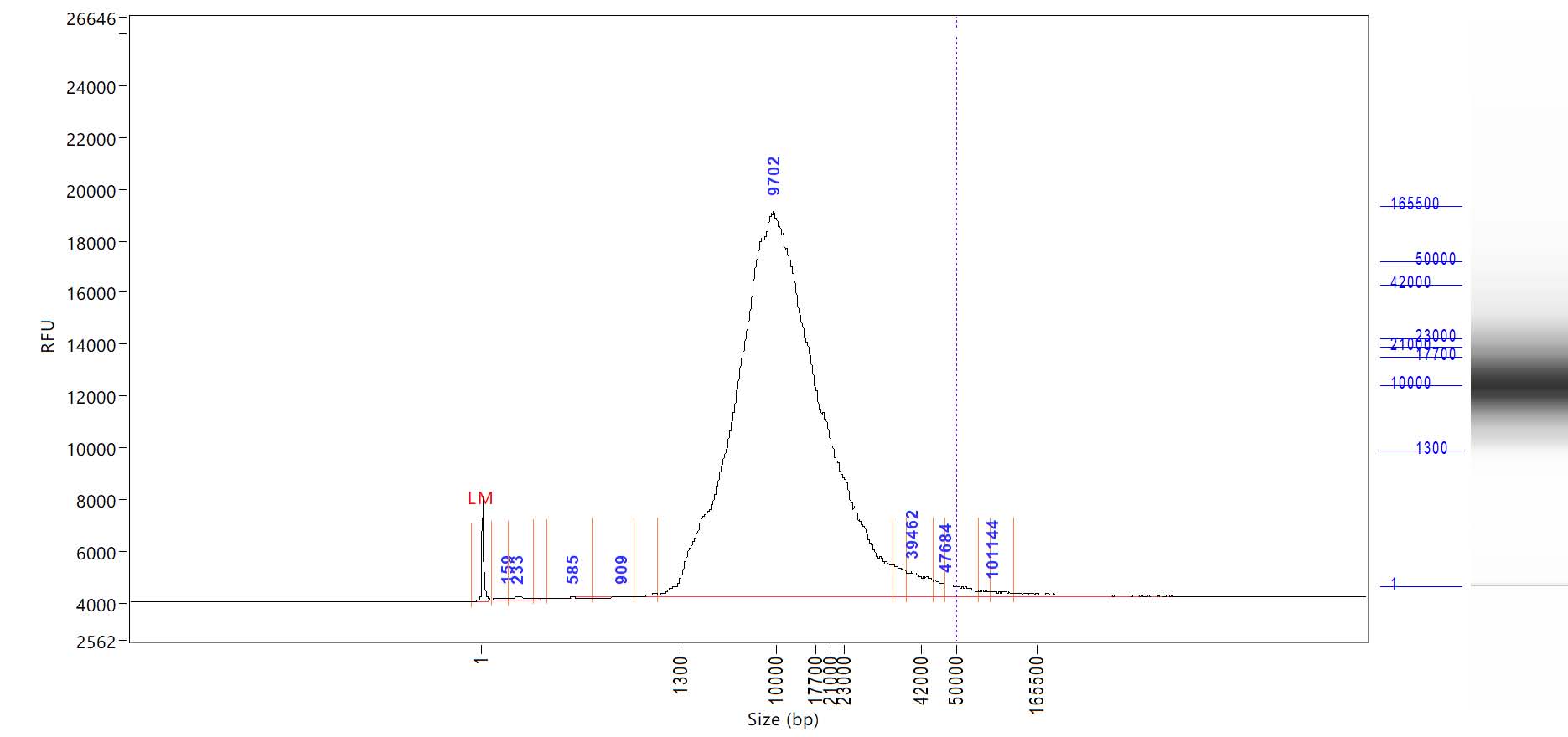

Distribution of fragment size (bp) and relative fluorescence units (RFU) for *Peromyscus attwateri.*

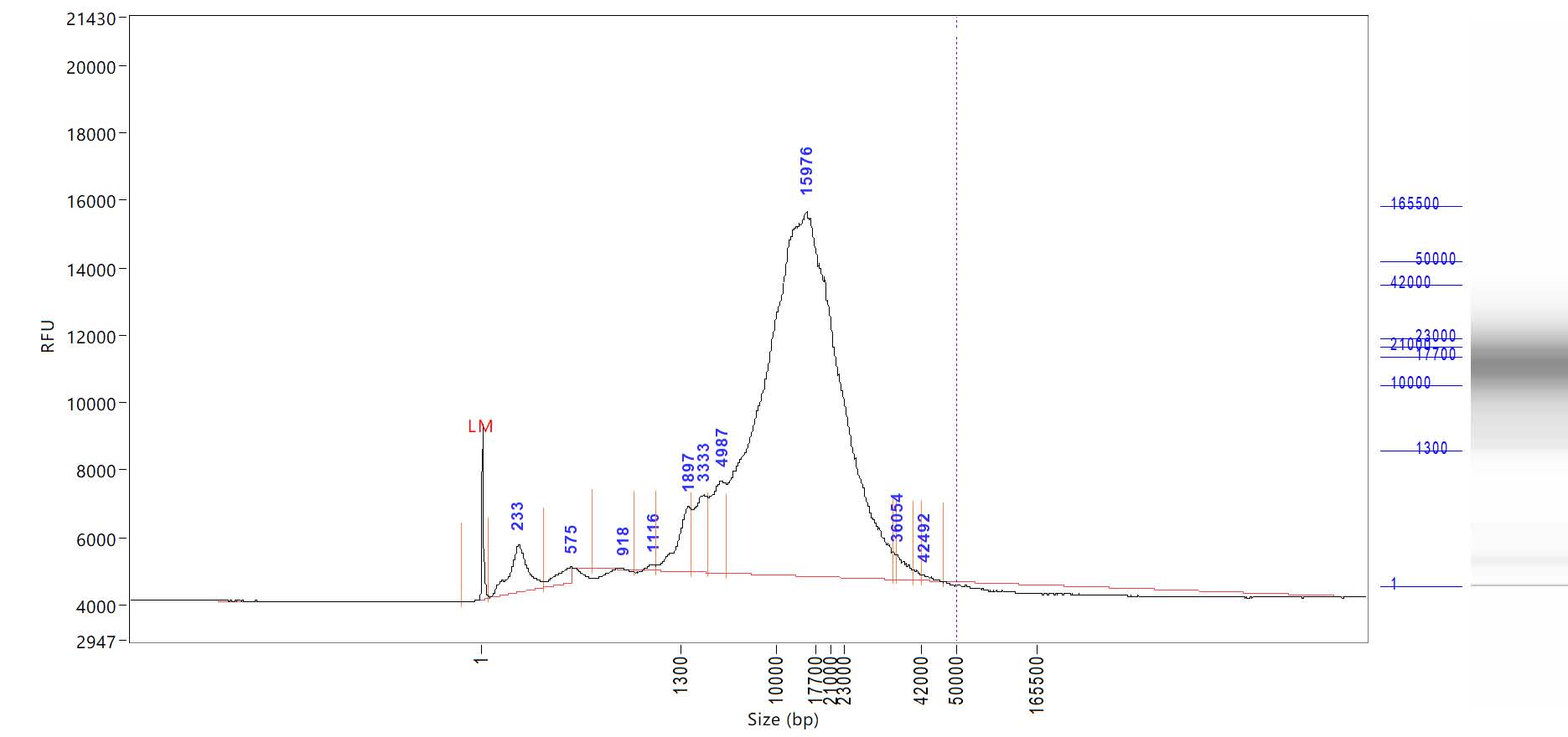
